# Meta Analysis of the *Ralstonia solanacearum* species complex (RSSC) based on comparative evolutionary genomics and reverse ecology

**DOI:** 10.1101/2021.12.05.471342

**Authors:** Parul Sharma, Marcela A. Johnson, Reza Mazloom, Caitilyn Allen, Lenwood S. Heath, Tiffany M. Lowe-Power, Boris A. Vinatzer

## Abstract

*Ralstonia solanacearum* species complex (RSSC) strains are bacteria that colonize plant xylem and cause vascular wilt diseases. However, individual strains vary in host range, optimal disease temperatures, and physiological traits. To increase our understanding of the evolution, diversity, and biology of the RSSC, we performed a meta-analysis of 100 representative RSSC genomes. These 100 RSSC genomes contain 4,940 genes on average, and a pangenome analysis found that there are 3,262 genes in the core genome (∼60% of the mean RSSC genome) with 13,128 genes in the extensive flexible genome. Although a core genome phylogenetic tree and a genome similarity matrix aligned with the previously named species (*R. solanacearum, R. pseudosolanacearum, R. syzygii*) and phylotypes (I-IV), these analyses also highlighted an unrecognized sub-clade of phylotype II. Additionally, we identified differences between phylotypes with respect to gene content and recombination rate, and we delineated population clusters based on the extent of horizontal gene transfer. Multiple analyses indicate that phylotype II is the most diverse phylotype, and it may thus represent the ancestral group of the RSSC. Additionally, we also used our genome-based framework to test whether the RSSC sequence variant (sequevar) taxonomy is a robust method to define within-species relationships of strains. The sequevar taxonomy is based on alignments of a single conserved gene (*egl*). Although sequevars in phylotype II describe monophyletic groups, the sequevar system breaks down in the highly recombinogenic phylotype I, which highlights the need for an improved cost-effective method for genotyping strains in phylotype I. Finally, we enabled quick and precise genome-based identification of newly sequenced Ralstonia strains by assigning Life Identification Numbers (LINs) to the 100 strains and by circumscribing the RSSC and its sub-groups in the LINbase Web service.

**IMPACT STATEMENT:** The *Ralstonia solanacearum* species complex (RSSC) includes dozens of economically important pathogens of many cultivated and wild plants. The extensive genetic and phenotypic diversity that exists within the RSSC has made it challenging to subdivide this group into meaningful subgroups with relevance to plant disease control and plant biosecurity. This study provides a solid genome-based framework for improved classification and identification of the RSSC by analyzing one hundred representative RSSC genome sequences with a suite of comparative evolutionary genomic tools. The results also lay the foundation for additional in-depth studies to gain further insights into evolution and biology of this heterogeneous complex of destructive plant pathogens.

**DATA SUMMARY:** The authors confirm that all raw data and code and protocols have been provided within the manuscript. All publicly available sequencing data used for analysis have been supplemented with accession numbers to access the data. The assembled genome of strain 19-3PR_UW348 was submitted to NCBI under Bioproject PRJNA775652 Biosample SAMN22612291. This Whole Genome Shotgun project has been deposited at GenBank under the accession JAJMMU000000000. The version described in this paper is version JAJMMU010000000.

## 1 INTRODUCTION

Named species generally correspond to groups of bacteria with pairwise genome similarity over a 95% average nucleotide identity (ANI) threshold and that also share a core set of phenotypes [1]. Bacterial plant pathogens rarely conform to this description. In contrast, many plant pathogenic bacteria belong to species complexes whose members share phenotypes but have pairwise ANI values below 95%. Further, one of the phenotypes that plant pathologists care most about, host range, varies widely among members of the same plant pathogen species.

The bacterial wilt pathogens in the *Ralstonia solanacearum* species complex (RSSC) are a notable example and the objects of this study. RSSC pathogens share a specialized habitat, the water-transporting xylem vessels and stem apoplasts of angiosperm plants, as well as a common pathology, lethal wilt symptoms [2]. Nonetheless, pairwise ANI of RSSC strains can be as low as 90.7%, and host ranges can vary dramatically between closely related strains that have pairwise ANI over 95% [3]. At the same time, many phylogenetically distant strains, with pairwise ANI below 95%, share host ranges [3, 4].

Genomic analyses place RSSC strains into four statistically supported phylogenetic clades that each share ANI values above 95% and correspond to geographic regions where the clades diversified [5]. These clades are known as phylotypes I, II, III, and IV with geographic origins in Asia, the Americas, Africa, and the Indonesian archipelago/Japan, respectively. Phylotype II can be further subdivided into IIA and IIB corresponding to two sub-clades [6]. Taxonomists formally divided the species complex into three species: *R. solanacearum*, corresponding to phylotype II; *R. pseudosolanacearum*, corresponding to both phylotypes I and III; and *R. syzygii*, corresponding to phylotype IV [7] (Figure 1).

**Figure 1.**
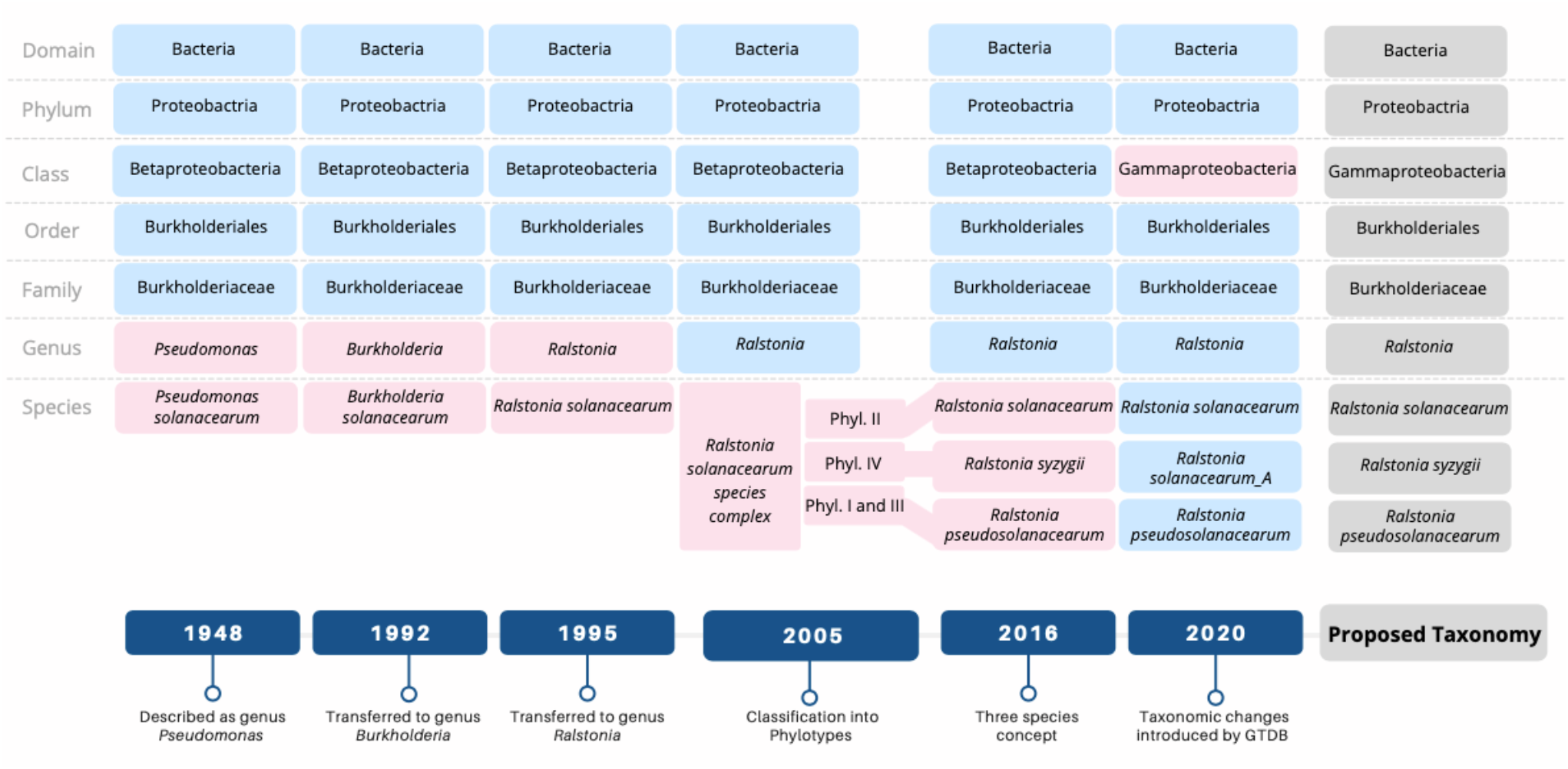
Major taxonomic revisions for the Ralstonia solanacearum species complex (RSSC). The bottom half depicts the timeline when these major changes were introduced, and the top half illustrates the predominant taxonomy used for each era. For each revision, pink boxes highlight changes to the classification, and blue boxes show levels that were unchanged. Taxonomic classification proposed through this paper is highlighted in grey color.

Describing the RSSC phylotypes as three named species conforms to taxonomic practice since RSSC clades are separated by genomic metrics and a few physiological traits correlate with the clades [5]. On the other hand, one could argue that there are not consistent differences in relevant pathogen behavior and ecology between clades to justify their division into separate species. Moreover, re-classification using new names leads to inconsistent naming of strains in the literature and in databases. The resulting confusion can interfere with one of the main goals of taxonomy: clear communication about organisms.

There is no simple resolution to this conflict. There are almost as many opinions about what a bacterial species is, and if bacterial species even exist, as there are taxonomists [8]. However, in today’s taxonomic practice, a pragmatic species “definition” is used. Bacterial species are commonly defined as groups of bacteria that have over 95% ANI to the name-bearing type strain of one species, have below 95% ANI to type strains of all other named species, and share a set of measurable phenotypes that distinguish them from members of other named species [1, 9]. Fortunately, genome sequence analysis now allows us to go far beyond ANI to infer many characteristics of groups of bacteria and to circumscribe bacterial species using a variety of species concepts, including the evolutionary, the ecological, and the pseudo-biological species concepts.

The evolutionary species concept considers species as independently evolving units [10]. Therefore, the investigation of evolutionary relationships or phylogenetics is the main approach for describing species based on this concept. The economic and technological accessibility of genome sequencing has allowed scientists to replace older approaches, such as DNA-DNA hybridization and 16S rRNA sequencing, with phylogenetic reconstructions based on whole genomes. Yet, even using all genes shared by a group of organisms may not precisely reflect their complete evolutionary relationships because of horizontal genetic exchange between sub-lineages [11]. However, it is hard to argue that there is anything that comes closer to representing evolutionary relationships than building a phylogenetic tree based on all gene sequences shared by the genomes under investigation, in other words, building a core genome phylogeny.

In a herculean effort, the genome taxonomy database (GTDB) team has built a phylogeny using protein sequences corresponding approximately to the core genome of all genome-sequenced prokaryotes [12, 13]. This effort has helped correct incongruencies in the taxonomic lineages of validly published species descriptions, which are often based on single gene 16S rRNA sequences. The names and lineages of these species descriptions can be found in the official List of Prokaryotic Names with Standing in Nomenclature (LPSN), which are reflected in large part by NCBI taxIDs [14]. Each time GTDB finds a genome that does not belong to a named species because it has a lower than 95% ANI to the type strain of a species, it creates a new species cluster with a placeholder name, e.g.: Escherichia coli_A. With respect to the RSSC, the GTDB changed a higher rank of the RSSC taxonomy: based on evolutionary distances inferred from genome sequences, the GTDB demoted the Betaproteobacteria to a subgroup nested within the class of the Gammaproteobacteria [13] (Figure 1). This shift in RSSC taxonomy was adopted by the microbial community profiling database SILVA with release 138 [15]. Importantly, GTDB does not resolve evolutionary relationships beyond the 95% ANI threshold (*i*.*e*., within species) since its goal is to improve “traditional” taxonomy based on the established ranks from kingdom to species and not to resolve evolutionary relationships within species.

The sequevar system was developed as a phylogeny-based taxonomy for within-species classification of the RSSC. This system coarsely estimates phylogenetic relationships of strains based on a multiple sequence alignment of a single DNA marker (a 750 bp region of the *egl* endoglucanase gene). Strains with similar sequences are assigned to <u>seque</u>nce <u>va</u>riant groups (sequevars) [16]. This can be considered a taxonomy focused on the “Evolutionary within-species concept”, with the expectation that some of the predicted relationships are inaccurate due to horizontal gene transfer (HGT). As the plant pathology community transitions from population genetics to population genomics, the ability of the sequevar system to estimate within-species phylogeny can be validated, which is one goal of this paper.

The ecological species concept defines a species as a group of bacteria that adapted to the same ecological niche [17]. Genomic comparisons can also provide insight into ecological species since bacterial adaptation necessarily involves a combination of gene gain/loss and allelic differentiation of gene sequences. For example, a pangenome analysis identifies gene families that are present or absent in different sets of genomes. These genome sets may represent groups that have adapted to different ecological niches and may thus represent different ecological species. Recently, the novel reverse ecology approach has gained traction [18]. This approach aims to identify populations that are in the process of adapting to an ecological niche based on frequent exchange of advantageous mutations during selective sweeps [19]. Putting this concept into practice, Arevalo and colleagues developed a tool, PopCOGenT, that assigns bacteria to distinct populations by identifying recent recombination events within sets of genomes and cessations of recombination between other sets of genomes [20]. Since the reverse ecology approach defines populations based on gene exchange, it also relates to the pseudo-biological species concept [21], which connects bacteria to the biological species concept, usually used for sexually reproducing eukaryotes. In the pseudo-biological species concept, gene exchange by homologous recombination during sexual crosses is replaced with gene exchange by HGT [22]. For example, the *Pseudomonas syringae* species complex has been proposed to represent a single species because HGT of virulence genes has been found to occur across the entire complex [23].

Because plant pathogenic bacteria with pairwise ANI values above 95% can have starkly distinct host ranges, plant pathologists have developed *ad hoc* within-species classification systems. In most pathogen groups, the “pathovar” concept is used to describe sub-species groups that cause the same disease on the same range of host plant species [24]. The “race” system is often used to describe strains within a pathovar that cause disease on different crop genotypes within the same species (for example in *Pseudomonas syringae* pv. *phaseolicola* [25]). The RSSC was never divided into pathovars, but for many years the term race was used in an attempt to divide strains by host range at the plant species level. This was never practically useful and eventually the RSSC race system broke down for two reasons. First, RFLP and sequence data revealed the “races’’ did not correspond to phylogenetic divisions [3, 26]. Second, most RSSC strains have very broad host ranges; it is not unusual for one strain to be able to cause disease on monocot and dicot hosts (e.g. banana and tomato [27] or potato and ginger [28]). As a result, most strains end up in a single unhelpful “Race 1” bin that includes members of all four phylotypes described above. In parallel, the RSSC was also subclassified into biovars based on *in vitro* physiological tests [29]. Once again, these biovars did not correspond to phylogenetic subgroups.

To alleviate the problem with the many different opinions about what should be considered a species, the confusion due to recurrent reclassification, and the various within-species classification schemes that are hard to use for non-specialists, we have developed a stable and neutral genome-based framework to circumscribe any of the above groups and to easily translate from one classification system to another. This system is based on genome similarity-based codes, called Life Identification Numbers (LINs) [30]. LINs consist of a series of positions with each position representing a different ANI threshold. ANI thresholds increase moving from the left to the right of a LIN. Therefore, bacteria with very low pairwise ANI do not share any LIN position (below 70% ANI). Bacteria with intermediate ANI (e.g. 95%), have identical LINs to an intermediate position (e.g., position F). Nearly identical bacteria (e.g., 99.99% ANI) have LINs that are identical up to, but not including, the rightmost LIN positions (e.g., position R or S). Therefore, LINs can precisely circumscribe any bacterial group with pairwise ANI values from 70% ANI, corresponding approximately to families and genera, to around 99.99%, corresponding approximately to clonal lineages. LINs have been implemented for numerous microbial genomes, including the representative genomes of GTDB, in the LINbase Web server [31].

The goal of this paper is to investigate RSSC classification through the lens of the different species concepts and within-species concepts by applying comparative evolutionary genomics and a reverse ecological approach to a set of representative, publicly available RSSC genomes. To translate this meta-analysis into applied utility, we then circumscribed the identified groups in the LINbase Web server, so that users can easily identify any new isolate based on its sequenced genome as a member of a named species, phylotype, population, or any other group within the RSSC.

## 2 MATERIALS AND METHODS

### 2.1 Selection of representative genomes

All publicly available genomes belonging to the three species (*Ralstonia solanacearum, Ralstonia pseudosolanacearum* and *Ralstonia syzygii*) were downloaded from the Assembly database of NCBI on September 5, 2020. Assembled genomes of strain *Ralstonia syzygii* R24 and Blood Disease Bacterium R229 were downloaded from the Microscope Microbial Genome Annotation and Analysis Platform - MaGe [32]. The genome of strain 19-3PR_UW348 was sequenced using the Pacbio Sequel II sequencing platform and assembled using Canu (version 2.0) [33]. It is included here as well (NCBI accession number JAJMMU000000000). All genome assemblies were assessed for quality using the CheckM (version 1.0.13) tool [34]. Genomes with completeness over 98%, contamination below 6%, number of contigs below 670, and N50 scores above 20,000 were retained. This genome set was further reduced by removing almost identical genomes to obtain a more even representation of the currently known genomic diversity of the RSSC. This was done using the LINflow tool (version 1.1.0.3) [35], retaining only one genome for each group of genomes that had reciprocal ANI values of over 99.975%. Preference was given to genomes of higher sequence quality and for which more published biological data were available.

### 2.2 Pangenome analysis and construction of the core-genome phylogenetic tree

The selected RSSC genomes were subjected to a pangenome analysis using PIRATE (version 1.0.4) [36]. To prepare the genome sequences for input to PIRATE, genomes were annotated using the PROKKA gene annotation tool (version 1.14.6) [37] with default settings. The annotated files were then used to obtain a core gene alignment whereby all genes present in at least 98% of the genomes were considered as core genes. The following parameters were used: `-a` to obtain a multiFASTA core gene alignment file as output and `-k` for faster homology searching with the `--diamond` option specified. The final core gene alignment file was used as input for IQtree (version 2.0.3) [38] using automated model selection to obtain a maximum-likelihood phylogenetic tree. The final phylogenetic tree was visualized using the ggtree [39] package in R. For the pangenome analysis, the PIRATE output file with all gene families was used to obtain the differences in gene content between different phylotypes. For phylotypes I and II, a gene was considered as a core phylotype gene if it was present in more than 95% of the genomes in a phylotype. Because of the much smaller number of genomes in phylotypes III and IV, presence in all but one genome was used as a rule. A score of 1 was assigned in case of gene presence and a score of 0 for gene absence. This assessment was performed for each gene in the pangenome for all 4 phylotypes (I,II,III,IV), resulting in a presence-absence matrix with genes as rows and phylotypes as columns (Supplementary Table 2). The matrix was then visualized through an upset analysis using the UpSetR [40] package in R.

### 2.3 ANI analysis

Pairwise average nucleotide identity (ANI) was measured for all representative genomes using pyani (version 0.2.10) [41] with default settings. The resulting matrix was used to construct a heatmap of ANI values using the function heatmap.2 under the gplots package [42] in R.

### 2.4 Recombination analysis

First, a recombination analysis of the RSSC was performed within the core genome. The core gene alignment and the phylogenetic tree obtained in the pangenome analysis were used as input to ClonalframeML (version 1.12) [43] with default parameters. The inferred recombination regions were used in two different analyses: (1) to find the genes in these regions using SAMtools (version 1.12) [44] with the command `intersect`; and (2) to build a recombination-free phylogenetic tree by masking the recombination regions using cfml-maskrc [45] and using the new recombination-free alignment as input to raxml-ng (version 1.0.3) [46] with the following parameters `--all -- model GTR+G --bs-trees 1000`. The tree was visualized using the ggtree [39] package in R.

Next, a recombination analysis was performed separately for each phylotype including the entire genome. For each phylotype, three different reference genomes (four for phylotype II; Table S3) were picked based on the CheckM results. The corresponding genomes were used as input to snippy (version 4.6.0) [47] to generate a whole genome SNP alignment mapped to each of the different reference genomes separately. The whole genome SNP alignment was used as input to gubbins (version 3.0.0) [48] to obtain the regions under recombination for each phylotype. The SAMtools `intersect` [44] function was used to find the genes in these regions.

### 2.5 Reverse ecology analysis

To obtain population predictions, inferred from the pairwise measurement of HGT, all of the representative genomes were used as input to PopCOGent (downloaded from https://github.com/philarevalo/PopCOGenT on March, 2021 [20].

### 2.6 Sequevar analysis

Automated sequevar assignments were generated using a custom bash script that takes a query genome sequence and compares it to a database of *egl* gene sequences (compiled by E. Wicker, CIRAD, France [49] using the command line version of Basic Local Alignment Search Tool: BLAST (version 2.9.0+) [50]. Sequevar assignment was made based on the best hit with 99-100% alignment, and results were cross-checked with data from the literature when available.

### 2.7 LIN assignment and LINgroup circumscriptions

All representative genomes and their metadata were uploaded into LINbase [31] for automated LIN assignment. LINgroups corresponding to groups identified here were circumscribed including a name, a description, and a link to this manuscript.

## 3 RESULTS and DISCUSSION

### 3.1 A core-genome phylogeny to determine evolutionary relationships

To classify the RSSC based on the evolutionary, ecological, and pseudo-biological species concepts, we needed to identify high quality genome sequences that best represent the described genetic diversity. We started with 167 publicly available genome sequences (Supplementary Table S1), from which we removed eleven low quality genomes that were fragmented into many contigs, had low genome completeness scores, had high contamination scores, or had a high number of ambiguous bases. From the remaining 156 genomes, we selected 100 genomes (Figure 2) best representing the known diversity of the species complex and limiting redundancy due to several nearly identical genomes present in the original set.

**Figure 2.**
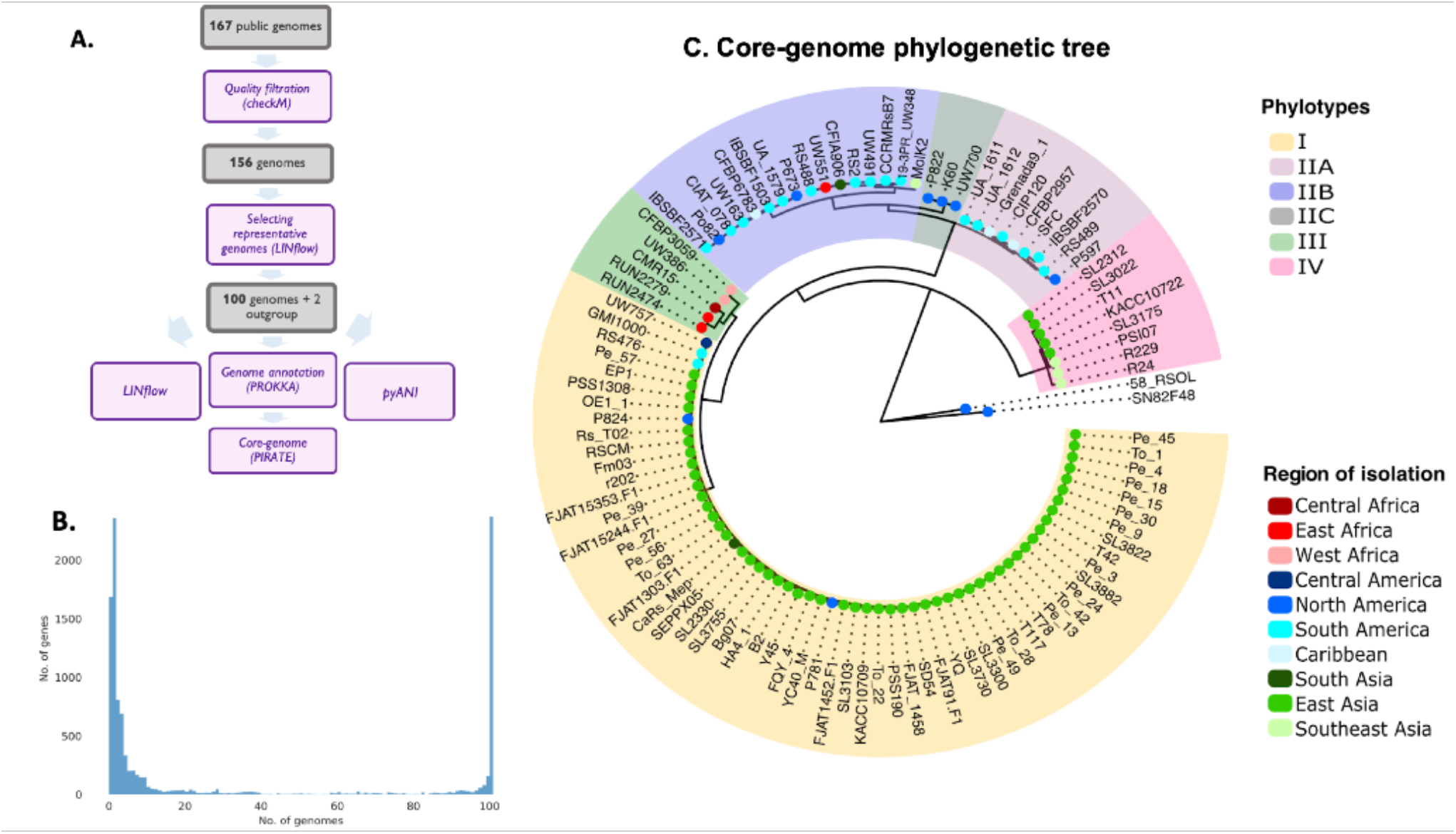
Core genome analysis for the representative genomes of RSSC. (A) Selection of the representative genomes. Purple boxes indicate the software used, and the grey boxes show the number of genomes left at each step. (B) The number of genomes that carry each gene in the pangenome. (C) Phylogenetic tree obtained with the core-genome analysis. All clades with high bootstrap values are included in the tree. Phylotypes of the strains are highlighted in different colors representing phylotypes I, IIA, IIB, III, IV. Based on the analysis, strains P822, K60, UW700 are classified as phylotype IIC. Colored dots at the node of each strain represent the region of isolation.

To uncover the phylogenetic relationships among the representative strains, we performed a pangenome analysis. This analysis revealed that 3,262 orthologous genes constitute the RSSC core genome (Table S2). A phylogenetic tree based on these core genes (Figure 2) clustered strains into clades corresponding to the four known phylotypes, with 59 strains belonging to phylotype I, 28 strains belonging to phylotype II (among which 9 and 16 strains belonged to phylotypes IIA and IIB, respectively, and 3 strains were intermediate between IIA and IIB), 5 strains belonging to phylotype III, and 8 strains belonging to phylotype IV. During this analysis, we identified one genome sequence that may be the result of a chimeric assembly between a phylotype I strain and a phylotype II strain: CRMRs218. This genome was published as a phylotype I strain [51], but in the core genome tree it formed a singleton branch basal to all phylotype II strains. Because of this ambiguity, the strain was excluded from further analysis.

Based on the geographic origin of strains, the phylogenetic tree is consistent with the hypothesis that the phylotypes diversified in different global regions [4, 52]. In fact, most phylotype I strains were isolated in continental Asia, phylotype II strains in the Americas, phylotype III strains in Africa, and phylotype IV strains in Indonesia and Pacific Islands (Figure 2). It is important to point out that the strains used here are not equally distributed between and within continents and thus neither are phylotypes. For example, strains belonging to phylotype III isolated in Africa are underrepresented (5% of total strains) compared to other phylotypes. East Asian strains represent 90% of the analyzed phylotype I strains, with most sequenced strains isolated in either South Korea or China. Although phylotype I is common in South Asia, only 1.7% of the sequenced phylotype I strains were isolated in South Asia. This uneven representation most likely reflects a bias in publicly available genome sequences from different geographic regions and is not a reflection of the actual geographic distribution and diversity of RSSC strains.

The phylotype II circumscription was consistent with the classification of strains based on the LPSN and GTDB classification systems of belonging to the named species *R. solanacearum*. Similarly, all phylotype I and III strains were consistent with the LPSN and GTDB classification of belonging to the recently named species *R. pseudosolanacearum* [7]. Phylotype IV strains correspond to *R. syzygii* as per LPSN taxonomy and “*R. solanacearum*_A” as per GTDB. It is important to note that many strains that are members of *R. syzygii* and *R. pseudosolanacearum* are listed as *R. solanacearum* in NCBI, because the genomes were submitted before the reclassification and adoption of the new species names by the scientific community.

### 3.2. Pangenome analyses provide a basis to investigate adaptation to ecological niches

One of the currently unanswered questions about the RSSC is to which degree the four phylotypes diverged from each other because of adaptation to different niches or because of allopatry. As a small step towards answering this question, we determined the congruences and differences in gene content between and within phylotypes.

Overall, the RSSC contained a total of 13,128 gene families, which represent the RSSC pangenome. The respective pangenome sizes of the individual phylotypes are: 4,023 (I), 3,329 (II), 3,909 (III), 3,971 (IV). An Upset plot was used to visualize the number of genes that are either shared by all strains of one phylotype and absent from all other phylotypes, *i*.*e*., the phylotype-specific core genes, or that are shared between subsets of phylotypes (Figure 3). Due to the above mentioned differences in the extent to which the genomic diversity within each phylotypes was sampled, it is difficult to make firm conclusions. Nonetheless, based on the available data, the core genome of phylotype II (3,329 genes) was considerably smaller than that of the other phylotypes (3,909-4,023 genes).

**Figure 3.**
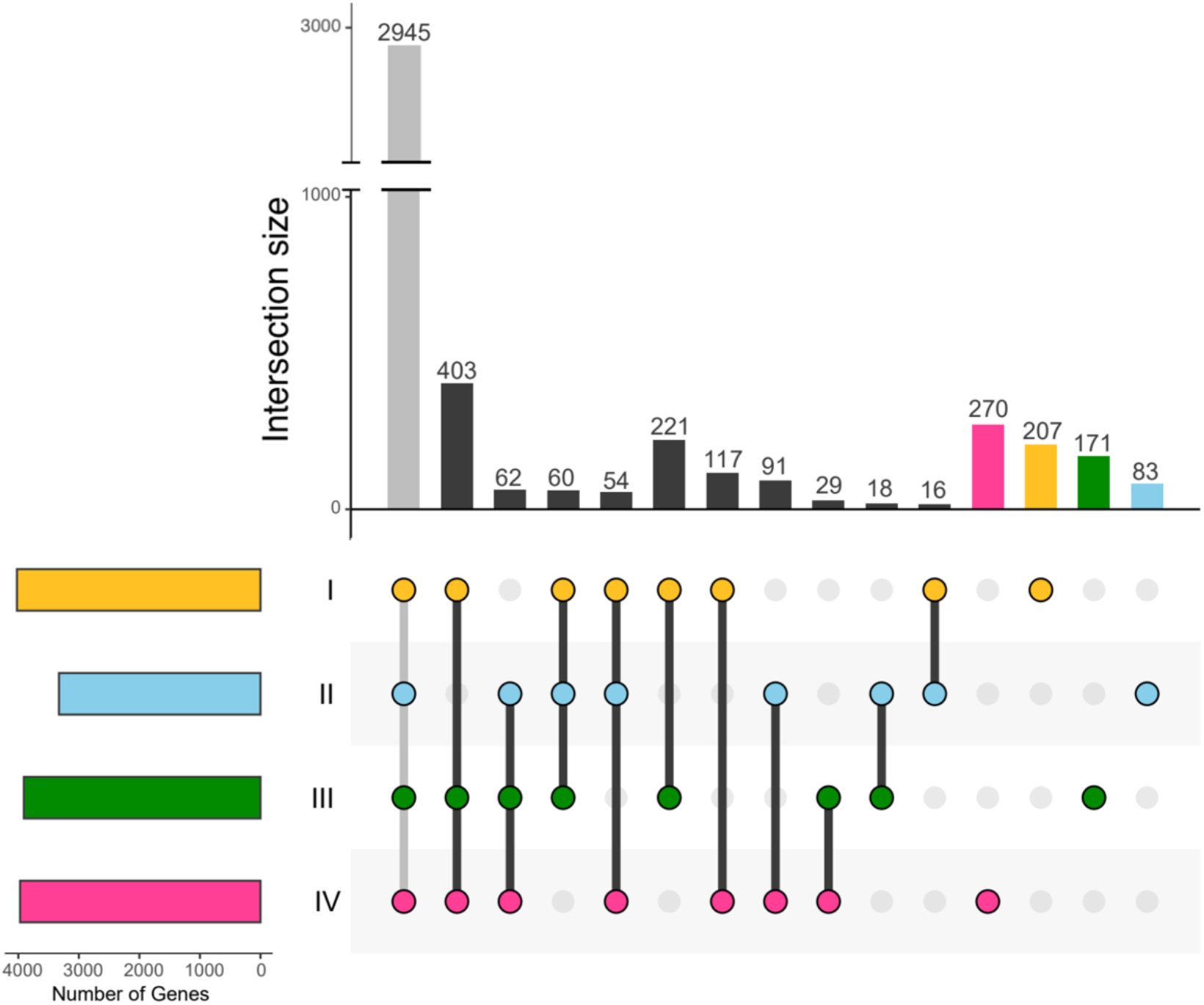
Pangenome analysis represented using an Upset plot to highlight how many genes are shared between phylotypes I, II, III, and IV. Each bar on the vertical bar chart represents the number of genes shared by the combination of phylotypes shown below the chart. The horizontal bar chart indicates the size of the phylotype-core genomes.

At the species level, *R. solanacearum* (phylotype II) has a core genome size (3,329 genes) very similar to the core genome size of *R. pseudosolanacearum* (phylotype I and III) (3,408). A surprising finding is the large core genome size of the *R. syzygii* species, which includes strains that cause the most phenotypically diverse diseases (Sumatra disease of cloves, banana blood disease, and classical bacterial wilts) [55]. However, the large size of the *R. syzygii* / phylotype IV core genome (3,971) may be an artefact due to the small number of phylotype IV genomes available.

When comparing gene content between phylotypes, phylotypes I and III share the most core genes with each other that are not core genes of the other phylotypes (221 genes). This is consistent with the shared membership of phylotypes I and III in the *R. pseudosolanacearum* species. Phylotypes I, III, and IV constitute the group of phylotypes that have the most genes in common that are absent from the core genome of the remaining phylotype, *i*.*e*., phylotype II in this case (403 genes). This is consistent with phylotype II having the smallest core genome and being the most diverse phylotype in regard to gene content.

### 3.3 ANI analysis confirms species boundaries and genome similarity-based clusters

After determining phylogenetic relationships and comparing gene content between strains providing the basis for investigating the RSSC from an evolutionary and ecological perspective, we calculated pairwise ANI between all 100 genomes (Figure 4 and Table S3). Since ANI is based on the average genetic distance of all DNA sequences shared between pairs of strains, it provides an orthogonal measure of genomic relationships beyond a core genome tree, which is limited to the genes shared by all 100 strains. In agreement with the core genome analysis, pairwise ANI clustered the genomes into the four phylotypes. Importantly, although phylotypes I and III formed distinct clusters, all strains in these two phylotypes had pairwise ANI values above 95%, which is consistent with these phylotypes being part of the same species.

**Figure 4.**
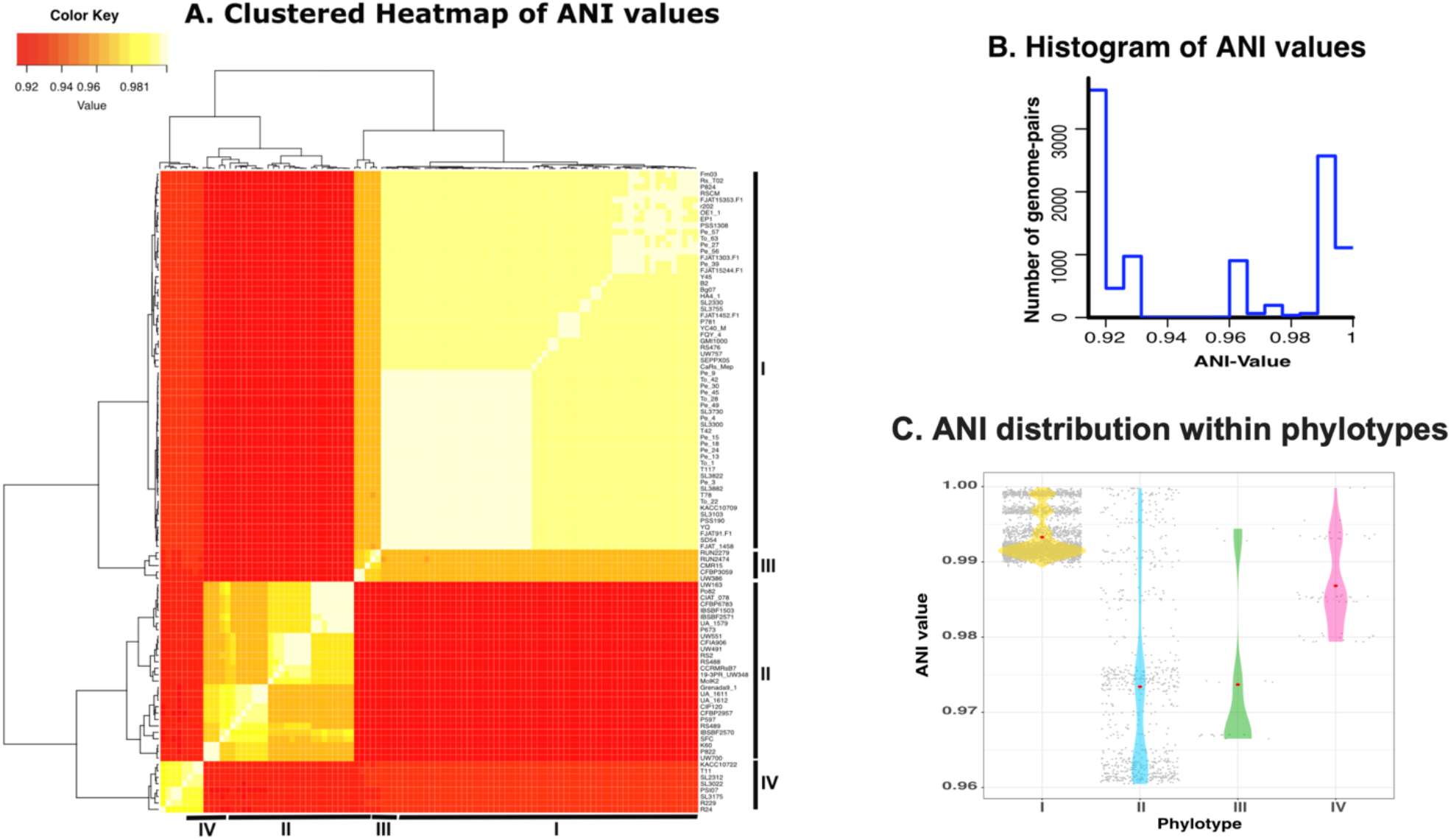
Average nucleotide identity (ANI) analysis for representative RSSC genomes. (A) Heatmap of pairwise ANI values for all genomes. (B) Histogram of pairwise ANI values among all paired genome combinations. (C) Pair-wise ANI distribution within each phylotype. Grey dots represent pairwise ANI between genomes belonging to the same phylotype, and red dots show the mean ANI for each phylotype.

Phylotype I strains had higher average pairwise ANI (99.35%) than other phylotypes (97.73% for phylotype II, 97.30% for phylotype III, and 98.67% for phylotype IV). Phylotype I appears to be the most genetically homogenous phylotype, but, as pointed out above, the genomic similarity could be an artefact stemming from the limited geographic distribution of most phylotype I genomes. If the high ANI among phylotype I strains is maintained as South Asian strains are sequenced, this may indicate that phylotype I emerged more recently in evolutionary time, possibly from within the wider genetic diversity of phylotype III.

Strains within phylotype II are characterized by relatively low ANI. Pairwise ANI indicates that there are three main subgroups. Strains in the sequevar 7 clade (K60, UW700, P822) had high pairwise ANI with each other (mean ANI 99.73%) and lower ANI with IIA and IIB strains (mean ANI 97.53% and 96.18%, respectively), which is consistent with sequevar 7 strains clustering as a phylotype separate from phylotypes IIA and IIB.

### 3.4 Recombination analyses provide a basis to identify biological and ecological species

Most RSSC strains are naturally transformable [56], and prior population genetics and genomics studies at the global, regional, and field scales have indicated that RSSC genomes are highly recombinogenic [52, 57–59]. To investigate whether the core genome phylogenetic tree was biased by recombination within the RSSC, we used ClonalFrameML to identify core genes that lack evidence of recombination. ClonalFrameML found recombination regions in 1,559 core genes (Table S4). The recombination regions detected by ClonalFrameML were masked and a recombination-free tree is shown in Figure 5B. While this tree maintained the main clades from the core genome tree shown in Figure 2 and 5A, the Southeast US clade (sequevar 7) shifted and became basal to phylotype IIA. This suggests that this clade’s basal-to-phylotype-IIB position in the core genome tree (Figure 2) could be due to recombination between its members and phylotype IIB strains rather than reflecting vertical inheritance.

**Figure 5.**
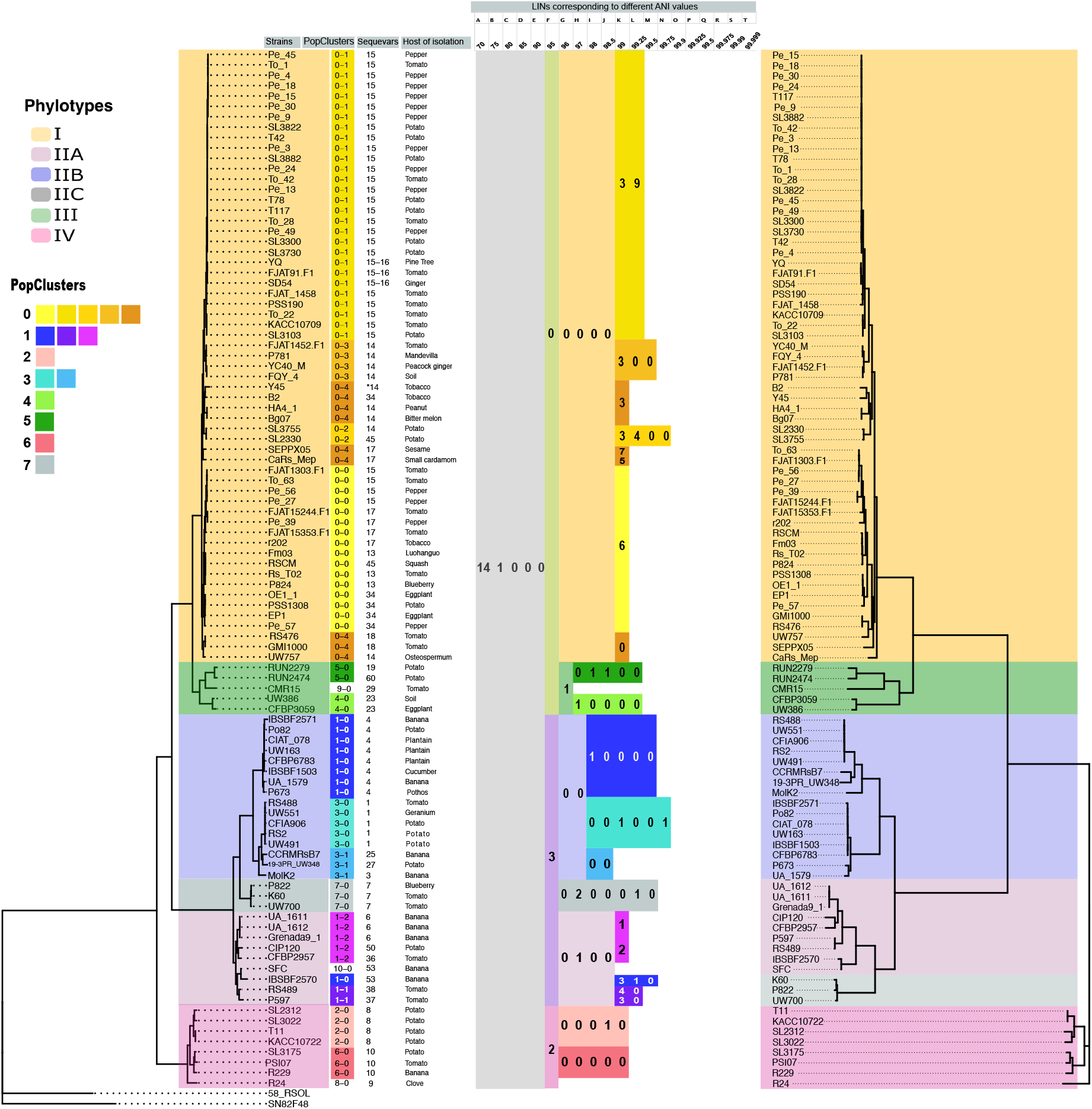
Comparison of core-genome tree, recombination-free tree, population clusters, sequevar types, and delineation of RSSC groups using LINs. The tree on the left is a vertical version of the core-genome phylogenetic tree from Figure 2. To the right of each strain name, assignments to population clusters, sequevars, and then the respective hosts of isolation. LINs corresponding to each group (the RSSC, named species, phylotypes, sub-phylotypes, and population clusters) are listed using colors matching each group. Newly sequenced genomes can be identified as members of these groups at www.linbase.org. A flipped recombination-free tree is depicted on the right.

Strains that have exchanged genes in recent history may belong to populations in the process of speciation, based on the ecological and biological species concepts. To determine which genomes belong to the same population based on recombination events in their entire genomes, we used PopCOGenT [20]. The population membership (“Pop Clusters”) of each genome is aligned to the core genome tree (Figure 5A). Most populations clustered phylogenetically related strains (18/20 PopClusters). In three cases, individual strains formed populations that only contain themselves (Phyl. III strain CMR15 in PopCluster 10-0, Phyl, IV strain R24 in PopCluster 11-0, and Phyl. II strain SFC in PopCluster 12-0), indicating that they may be the only sequenced members of under-sampled populations. However, there were two PopCOGenT clusters that were polyphyletic: PopCluster 2-0 contained 8 IIB-4 strains and a IIA-57 strain IBSBF2570, while PopCluster 0-4 contained 8 phylotype I strains from three distinct branches on the core genome tree.

Genes that are frequently transmitted horizontally between strains may play a role in adaptation (the ecological species concept). Therefore, in addition to PopCOGenT, we ran the independent recombination tool Gubbins [48] to detect recombination in the RSSC using 13 reference genomes (3 genomes for phylotype I, III, and IV and 4 genomes for phylotype II). The results are summarized in Figure 6. Table S4 contains the estimated number of recombinations for each gene in the 13 reference genomes. As expected, mobile genetic elements (transposases, integrases, and phage associated proteins) were highly recombinogenic genes. Many of the highly recombining genes are type III secreted effectors, which RSSC strains use to manipulate plant host physiology and immunity. The high plasticity of type III effector repertoires is well known in RSSC strains [60]. Additionally, glycoside hydrolases, polygalacturonases, and endoglucanases displayed evidence of frequent recombination. Endoglucanases are involved in adaptation of *Xanthomonas* spp. to vascular vs. apoplastic niches [61], but variation in plant cell wall degrading enzyme repertoires has not been investigated for RSSC. Several classes of genes involved in inter-microbial interactions were recombinogenic: non-ribosomal peptide synthetases and polyketide synthases [62], type VI secretion system genes like Vgr, PAAR, and putative effector/immunity pairs [63], and hemagglutinin-like proteins that are hypothesized to be contact-dependent inhibition (CDI) systems in RSSC [59]. Investigating the functional diversity of the recombining genes may shed light on how interactions with plant hosts, microbial competitors, and novel abiotic environments shape the evolution of RSSC lineages.

**Figure 6.**
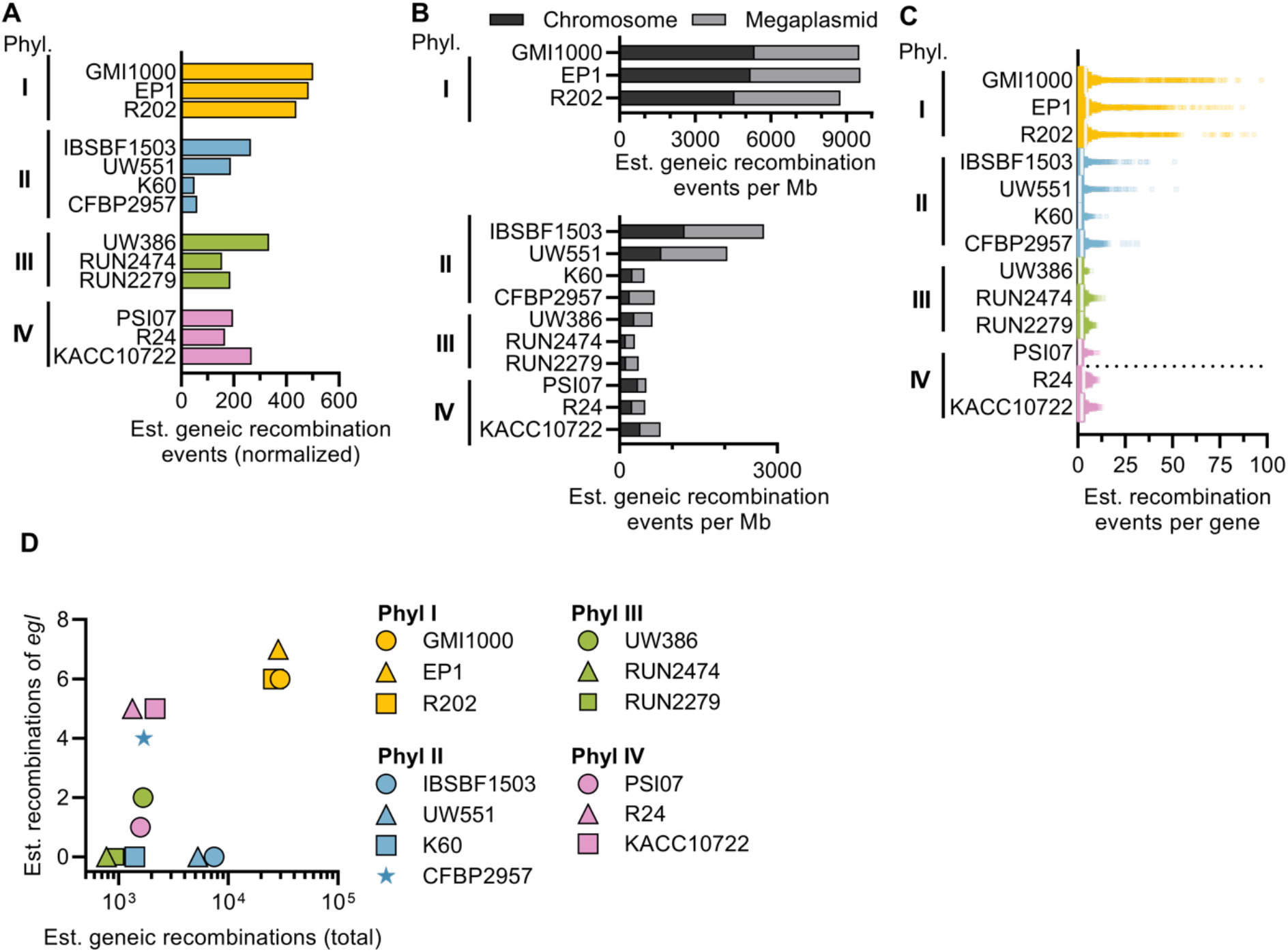
Comparison of estimated recombination for representative RSSC genomes from each phylotype. Genes with putative recombination events were identified using Gubbins [48]. (A) The number of recombination events for each genome, normalized by the number of genomes of each phylotype in the genome set. (B) The number of recombination events on the chromosome vs. megaplasmid, normalized by the length of the replicon. (C) Estimated number of recombination events detected for each gene (dots). (D) Comparison of the number of recombination events for the sequevar marker gene (*egl*) vs. the total recombination events for each genome.

### 3.5 Speculation on the relative evolutionary ages of phylotypes

Overall, our comparative genomics analyses suggest that either phylotype II (*R. solanacearum*) or phylotype IV is the most ancestral phylotype within the RSSC. Phylotype II genomes have the lowest average pairwise ANI value and phylotype II has the smallest core genome. Their lower recombination rate is also in line with higher sequence diversity since higher sequence diversity decreases the success of homologous recombination. All these results suggest that phylotype II is more diverse compared to the other phylotypes and, thus, could have emerged first. These findings are also consistent with an earlier study in which 29 RSSC genomes and 73 MALDI proteomes were compared [5]. Surprisingly though, phylotype IV is on the most basal branch in the core genome tree (Figure 2), as it was in a previous multi-locus sequence analysis tree [52]. This suggests that phylotype IV is the most ancestral phylotype. This inconsistency could be due to uneven sampling among phylotypes. The genomic diversity in phylotype IV may be under-sampled, and if additional genomes of diverse phylotype IV strains were to be sequenced, it might become more diverse than phylotype II. On the other hand, the basal position of phylotype IV might have been influenced by the choice of outgroup strains. If phylotype IV strains acquired genes from environmental *Ralstonia* closely related to the chosen outgroup strains, recombination could make phylotype IV seem more closely related to the outgroup strains than they are by vertical inheritance. Therefore, we cannot firmly conclude which phylotype is most ancestral based on available data. On the other hand, there is one clear interpretation about relative ages of the phylotypes. Phylotype I is the least diverse phylotype that also branches off the latest as a lineage from phylotype III, making it likely the phylotype that most recently emerged and expanded.

### 3.6 Comparing sequevars (*egl* trees) with the core genome phylogeny and populations

The global plant pathology community has widely adopted the sequevar taxonomic system to classify *Ralstonia* strains at the within-species level. Over 5,000 strains from over 88 regions have been assigned to over 70 sequevar groups [64]. Because the sequevar system is based on a single genetic marker (750 bp of the *egl* gene), and RSSC genomes often recombine, we predicted that the *egl* gene may have recombined between strains. Indeed, *egl* recombination events were detected in 3 of 3 phylotype I, 1 of 4 phylotype II, 1 of 3 phylotype III, and 3 of 3 phylotype IV reference genomes used in the Gubbins analysis (Fig 6D and Table S4). We and other plant pathologists have deposited over 4,500 “(egl) gene, partial cds” sequences of RSSC isolates to the NCBI nucleotide database, but our results suggest that recombination of *egl* within the RSSC may limit the sequevar taxonomy’s ability to accurately estimate phylogenetic relationships.

With evidence that *egl* may be horizontally transmitted between RSSC strains, we investigated whether the sequevar system and trees constructed with *egl* sequences reflect phylogenetic relationships of strains. We extracted the partial *egl* nucleotide sequences from each of the 100 RSSC genomes and aligned them with reference sequences to assign sequevars to each genome (Table S1 and Figure 5A). The sequevar assignments were monophyletic in the tested genomes for phylotype II (28 genomes assigned to 12 sequevars), III (5 genomes assigned to 4 sequevars), and IV (8 genomes assigned to 3 sequevars). Sequevar I-18 and sequevar I-13 mapped to single branches of the tree, so these sequevars may be monophyletic. However, most of the phylotype I sequevars were highly polyphyletic. Five of the phylotype I sequevars (I-14, I-15, I-17, I-34, and I-45) were assigned to distinct branches within the phylotype I.

Overall, our results and prior work [57] indicate that the sequevar system is not informative for describing within-species relationships for phylotype I RSSC. The polyphyletic phylotype I sequevars are probably due to the inter-related phenomena of phylotype I’s low genetic diversity and higher recombination. This suggests that improved methods for classifying within-species groups of phylotype I are needed, and PCR assays that target insertions/deletions might be a cost-effective method to prioritize strains for whole-genome sequencing [54]. On the other hand, the sequevar system appears to robustly reflect phylogenetic relationships for the diverse phylotype II strains. As more phylotype III and phylotype IV genomes become available, it will be useful to test whether the sequevar system works well in these phylotypes.

### 3.7. Using LINs to circumscribe RSSC groups for easy genome-based identification

In the LIN system, genomes are classified based on genome similarity without deciding on any *a priori* group boundaries. LINs can thus be used to circumscribe species complexes, species, or within-species groups and place any genome within these groups. If the breadth of a taxon is defined based on an ANI distance from the type strain, this can be done based on the LIN assigned to the type strain. For example, K60 is the type strain of *R. solanacearum*, and the LIN of K60 up to the F position (corresponding to 95% ANI) in the LINbase web server is 14_A_1_B_0_C_0_D_0_E_3_F_. Therefore, the LIN of the *R. solanacearum* species is 14_A_1_B_0_C_0_D_0_E_3_F,_ and each genome that has the same LIN at these positions can be immediately identified as a member of the species *R. solanacearum*. As shown in Figure 6, the LIN for *R. pseudosolanacearum* is 14_A_1_B_0_C_0_D_0_E_0_F,_ and the LIN for *R. syzygii* is 14_A_1_B_0_C_0_D_0_E_2_F_.

If a type strain genome is not available for a group or a group does not have a predetermined ANI breadth (because it is not a species), the group can still be circumscribed based on the LIN positions shared by its members. Since we added the 100 RSSC genomes used in this study to the LINbase web server and assigned LINs to each of them, we were also able to circumscribe the RSSC and its phylotypes, sub phylotypes, and population clusters so that any newly sequenced genome can be identified not only as a member of a species but also as a member of any of these other groups. In Figure 6, we report the LINs corresponding to each of these groups. While the LINs assigned to each individual genome are not shown in the figure, they are stored in Table S1 and in LINbase and can be used to circumscribe even more highly resolved groups corresponding to individual genetic lineages within the RSSC. Whole genome-based LINs could thus be used to replace the single marker gene-based sequevar system, which we have shown to contradict core genome phylogeny for phylotype I.

## 4.0 Conclusion

In conclusion, we have shown how a genomic meta-analysis can be used to classify the RSSC according to the evolutionary, biological, and ecological species concepts. We circumscribed validly published named species, phylotypes, clades within phylotypes, sequevars (when possible), and populations. We determined how extensively genes are shared within and between phylotypes and which genes most frequently recombine. We also provided the basis for further, more in depth, investigations of the RSSC. LINbase makes it straightforward to circumscribe any additional groups based on additional sampling and genome sequencing of the diversity within the RSSC and additional genomic comparisons and phenotypic tests. Any new isolate with a draft genome sequence can then be precisely identified as a member of any of these groups to help inform basic research, disease management, and biosecurity regulations.

## Acknowledgements

We thank our colleague Emmanuel Wicker (CIRAD, France) for providing reference egl sequences and Noah A. Kinscherf, Jessica L. Prom, and Alicia N. Truchon for genomic DNA extractions at UW-Madison. Additionally, we thank Stéphane Poussier (University of La Réunion) and Jonathan Jacobs (The Ohio State University) for helpful discussions about sequevar typing of phylotype I strains.

